# Aging exacerbates muscle loss induced by semaglutide treatment in female mice

**DOI:** 10.64898/2026.11.11.766063

**Authors:** Subhasmita Rout, Takuya Karasawa, Elizabeth R. Moore, Ran Hee Choi, Shinya Watanabe, Amandine Chaix, Micah J. Drummond, Katsuhiko Funai

## Abstract

**Background:** Glucagon-like peptide-1 receptor agonists (GLP-1RAs), including semaglutide, have transformed the pharmacological treatment of obesity. While highly efficacious in promoting weight loss, these drugs also induce substantial reductions in lean mass, raising concerns for skeletal muscle loss, particularly in older individuals vulnerable to sarcopenia. Nevertheless, there are currently no published human studies that investigated the effects of GLP-1RAs on skeletal muscle mass using gold standard methods. Meanwhile, studies in young mice demonstrate that skeletal muscle loss induced by GLP-1RAs might not be a major concern. The purpose of the current study was to investigate the effects of semaglutide on skeletal muscle mass and strength in aged mice.

**Methods:** Eighteen-month-old male and female C57BL/6J mice were fed a high-fat diet (HFD) for 12 weeks to induce obesity. At 21 months of age, mice received daily subcutaneous injections of vehicle or semaglutide (3 or 30 nmol/kg) for 3 weeks. Body composition was assessed weekly by nuclear magnetic resonance (NMR) spectroscopy. After 3 wks, mice were terminated to assess masses of individual organs including skeletal muscle. Skeletal muscle force-generating capacity was assessed by ex vivo electrical stimulation-induced contraction. Myofiber cross-sectional area and fiber-type composition were also assessed.

**Results:** Semaglutide induced dose-dependent weight loss in both sexes, with reductions of 11.3% and 20.2% in males and 21.1% and 26.1% in females receiving 3 or 30 nmol/kg dose, respectively. There was a substantial sex-difference in the loss of lean mass, with relatively minor reductions in males (2.3% and 6.6%) and greater loss in females (8.6% and 8.3%). Skeletal muscle mass was largely preserved in males but females exhibited greater muscle mass loss, including 14.2% reduction in quadriceps mass at 30 nmol/kg of semaglutide treatment. In contrast, skeletal muscle force-generating capacity was not compromised in semaglutide-treated females, while males exhibited 19.2% reduction in force generating capacity of the extensor digitorum longus (EDL) muscles at the 30 nmol/kg dose.

**Conclusion:** Unlike young mice, semaglutide treatment may induce meaningful loss in skeletal muscle mass in aged female mice and muscle strength in aged male mice. These findings highlight the importance of considering age as a potential factor that may negatively influence muscle mass and function in humans treated with GLP-1RAs. Clinical studies examining the influence of pharmacological weight loss on skeletal muscle mass and strength should consider both age and sex as a potentially important determinant of these outcomes.

## Introduction

Glucagon-like peptide-1 (GLP1) receptor agonists and other incretin mimetics have revolutionized obesity medicine with unprecedented efficacy for clinically meaningful weight loss and resolving obesity-associated metabolic complications [1–3]. In contrast, there has been reports that these drugs promote disproportionately greater loss of lean mass, raising concerns that they might represent loss of functional skeletal muscle mass that causes weakness [1, 4]. Nevertheless, lean mass quantified by DEXA are known not to accurately reflect changes in skeletal muscle mass, raising the question on the effect of pharmacological weight loss on the loss of skeletal muscle mass [5]. Multiple groups are currently conducting clinical studies utilizing gold-standard measurements for skeletal muscle mass (magnetic resonance imaging or D3-creatinine dilution) to study the effect of incretin mimetics on muscle mass. Meanwhile, rodent models present opportunities to directly assess skeletal muscle mass more quickly and invasively than they can be performed in humans. In our previously published study, we demonstrated that semaglutide-induced weight loss did not produce meaningful reductions in skeletal muscle mass in young male mice, despite substantial reductions in body weight and fat mass [6]. These findings have been supported by some preclinical studies [6–8], while others suggest that pharmacological weight loss may induce meaningful muscle loss [9–11].

Obesity remains highly prevalent (38.9%) among older adults above 60 years of age [12]. Thus, GLP-1RAs are increasingly prescribed to older adults, a population that stands to benefit considerably from effective weight reduction. Nevertheless, concerns regarding the potential muscle loss in these patients are also considerably higher, given their susceptibility to age-related skeletal muscle loss and functional decline. Furthermore, not only is aging associated with muscle atrophy and weakness, it is also associated with impaired recovery from muscle loss [13]. Considering the increased susceptibility for muscle loss, the concern for muscle loss during pharmacologic weight loss is also substantially higher in older individuals.

In response to these concerns, we re-examined the effects of semaglutide-induced weight loss on skeletal muscle mass and force-generating capacity in aged mice. We also report the data in both males and females. While young female mice are resistant to diet-induced weight gain and thus not ideal for weight loss studies, older female mice gain weight similarly to older male mice. These phenotypes allowed us to examine the effects of semaglutide-induced muscle loss in both aged male and female mice. Our findings demonstrate a somewhat concerning magnitude of loss in skeletal muscle mass in aged female mice and loss in force-generating capacity in aged male mice.

## Methods

### Animal Model and Experimental Design

Seventeen-month-old male and female mice C57BL/6J were acquired from the National Institute on Aging (NIA). Upon arrival, mice were housed in a temperature-controlled room (22°C) with a 12-hour light/dark cycle and provided with a standard chow diet and deionized water ad libitum. Mice were allowed to acclimate for a period of 4 weeks. At eighteen-month mice were fed 42% High-fat-diet (HFD) (Cat# TD.88137) till end of the study and body mass and food consumption were measured weekly till 12 weeks of HFD (before starting treatment). The body composition was measurement by nuclear magnetic resonance (NMR) before starting HFD and semaglutide treatment. At twenty-one-month, mice were randomly divided into two experimental groups where one group received daily subcutaneous injections of vehicle (phosphate-buffered saline, PBS) or 3 nmol/kg body mass or 30 nmol/kg body mass semaglutide (Novo Nordisk). Another age matched lean control group of mice maintained on standard chow diet throughout the experiment period. During the treatment period, body mass and food consumption were measured daily and body composition were measured weekly. After 3 weeks of treatment, before the termination the mice were fasted for 2 hours followed by intraperitoneal injection of 80 mg/kg ketamine and 10 mg/kg xylazine to proceed for the dissection. The muscle tissues, including tibialis anterior (TA), extensor digitorum longus (EDL), plantaris (PLA), soleus, gastrocnemius, and quadricep were harvested, snap-frozen in liquid nitrogen, and stored at -80°C for subsequent biochemical and molecular analyses. All animal experiments were performed with the approval of the Institutional Animal Care and Use Committees at the University of Utah.

### Body Composition

Whole-body fat and lean mass were determined using quantitative nuclear magnetic resonance (NMR) spectroscopy (Minispec, Bruker) on 17 months of age, 18 months of age and 21 months of age just before the treatment was started. The body composition was again measured after 3 weeks of post treatment.

### Skeletal Muscle Force-generating Capacity

Muscle force capacity was assessed using an ex vivo system (Model 801C, Aurora Scientific, Aurora, Canada). The EDL and soleus muscles were freshly harvested and then securely fastened at their tendons. Each muscle was individually mounted within a tissue bath, attached to an anchor at one end and a force transducer at the other. The bath contained continuously oxygenated with Krebs-Henseleit buffer or KHB (116 mM NaCl, 4.6 mM KCl, 1.2 mM KH_2_PO_4_, 25.3 mM NaH_2_CO_3_, 2.5 mM CaCl_2_, 1.6 Mm MgSO_4_) and maintained at 37°C. Once the optimal length was established, muscles were incubated in fresh KHB for a 5-minute equilibration period, following this, force-frequency was initiated. Muscles were stimulated at progressively increasing frequencies (10, 20, 30, 40, 60, 80, 100, 125, 150, and 200 Hz), with a 2-minute rest interval between each stimulation frequency. After the force measurements, muscle length and mass were precisely measured to calculate the cross-sectional area (CSA). Force production data were analyzed using Aurora Scientific’s DMAv5.321 software. Results are reported as specific force, which is calculated by normalizing the absolute force to the muscle’s CSA.

### Tissue Sectioning and Histological Analyses

Freshly harvested TA muscles were embedded in optimal cutting temperature media and were snap freeze in liquid nitrogen immediately and stored in -80°C freezer. Muscle tissues were sectioned at 8 μm thickness with the Leica CM1860 cryostat (Leica Biosystems) where the Cryostat chamber was maintained with a temperature of -20°C during the period. The tissue sections were collected on a positively charged coated slide and stored in -20°C for histology. To determine the fiber type in the TA muscle, the muscle sections were retrieved from the -20°C and allowed to incubate at room temperature for 15 min. Muscle sections were prefixed with 10% paraformaldehyde for 10 minutes followed by 1X PBS washing. Then sections were incubated with Mouse-on-Mouse (M.O.M) blocking reagent for 1 hour at room temperature (RT) to block the endogenous immunoglobulins following overnight at 4°C incubation with specific primary antibodies targeting different myosin heavy chain (MHC) isoforms: MHC I (DHSB, BA-D5), MHC IIa (DHSB, SC-71) and MHC IIb (DHSB, BF-F3) with 2.5% goat serum in 1X PBS. On the next day, muscle tissue sections were incubated with anti-mouse secondary antibody IgG2b (A 21242), IgG1 (A 21120) and IgM (A 21426) and wheat germ agglutinin (WGA) Alexa Fluor 488 conjugate (Thermo Fisher Scientific, 0.004 mg/mL) with 2.5% goat serum in 1X PBS for 1 hour at RT followed by PBS wash. Muscle sections were mounted with mounting medium (Vector Laboratories, H-1000) and imaged using Zeiss Axioscan 7 Slide Scanner (Carl Zeiss). The muscle fiber typing was analyzed using cellpose-SAM model as previously described.

### Statistical Analysis

GraphPad Prism 10.4.1 software was used to do statistical analysis such as ordinary one-way ANOVA to compare the terminal body weight, and lean tissue difference and two ways of variance (ANOVA) to analyze the body weight, food consumption and muscle force in semaglutide treated and vehicle groups in both male and female mice. All data are represented as mean ± SEM, and statistical significance was set at p ≤ 0.05.

## Results and Discussion

### Semaglutide Induces Dose-Dependent Weight Loss in Aged Male and Female Mice

Group-housed 18 months-old male and female C57BL/6J mice were fed a high-fat diet (HFD, 42% calories from fat, Inotiv: TD.88137) for 12 weeks to establish diet-induced obesity and were subsequently randomized at 21 months of age to receive daily subcutaneous injections of vehicle or semaglutide for 3 weeks while remaining on HFD (Figure 1A). An age-matched lean control group was maintained on chow diet throughout the experimental period. Similar to our previous study in young diet-induced obese mice, 3 or 30 nmol/kg dose of semaglutide were tested [6]. We chose the 18-21 months old mice because C57B/6J mice spontaneously begin to lose adiposity around 22 months of age, and we wanted to study semaglutide-induced weight loss independent of this effect. It’s important to note that the onset of sarcopenia in this strain of mice is around 26-28 months old, so skeletal muscle masses of mice at this age are equivalent to those in adult mice. While not directly comparable, 18-21 months old mice are roughly equivalent to humans in their 60s [14], which make up a substantial portion of individuals that are on incretin mimetics, and the age when functional decline begins to more directly influence quality of life [15].

**Figure 1.**
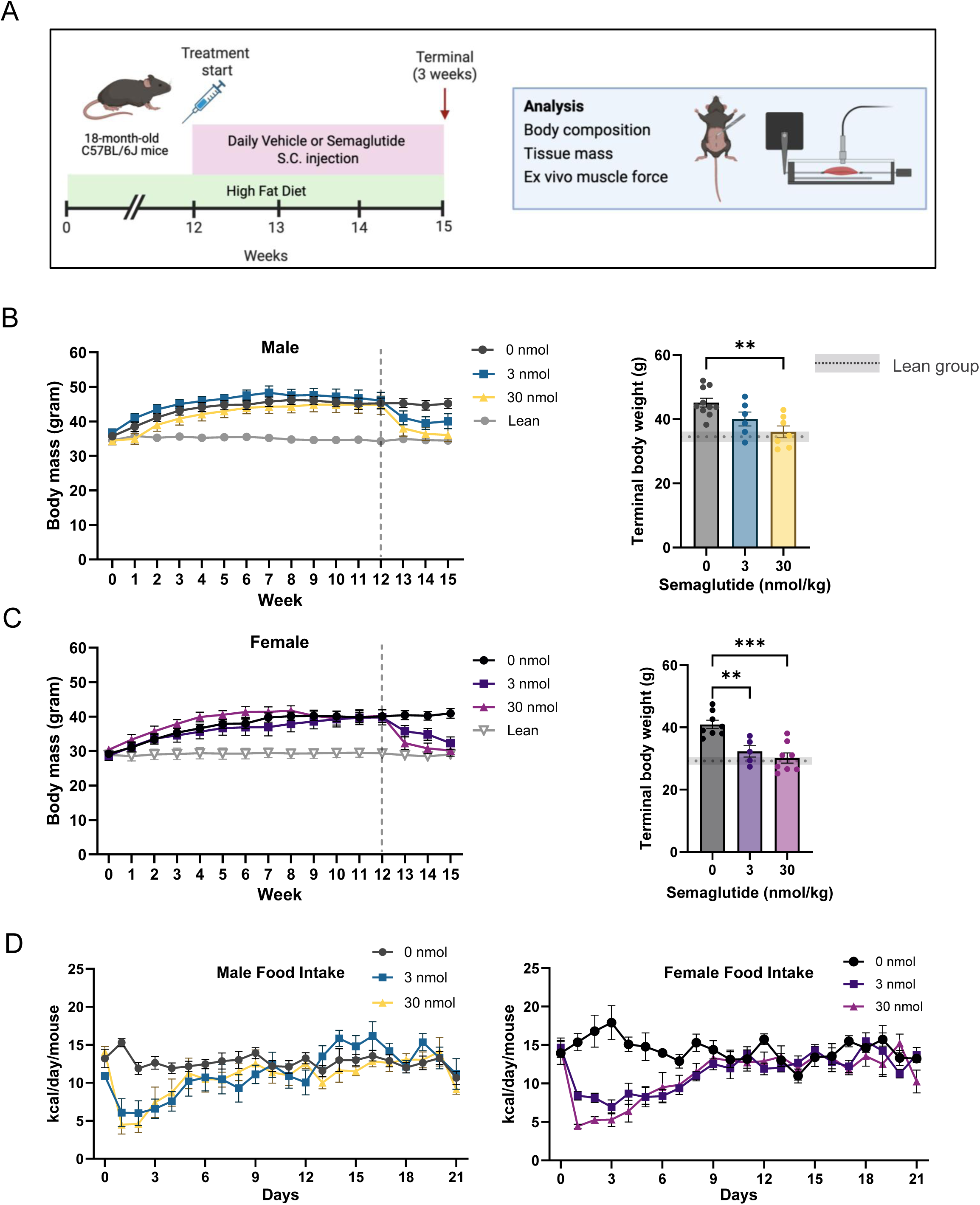
Effects of semaglutide treatment on body mass and food intake in aged male and female mice. (A) An experimental schematic. (B) Body mass progression throughout the experimental period and the terminal body mass in male mice. (C) Progression of body mass throughout the experimental period and the terminal body mass in female mice. (D) Food intake during semaglutide treatment in both male and female mice. All Data is represented as mean ± SEM. * p<0.05, ** p<0.01, *** and p<0.001, significant difference vs. corresponding controls. n = 6-10/group for male and n = 5-8/group for female mice.

All mass data from 0 (vehicle), 3, or 30 nmol/kg semaglutide-treated mice are shown in Table 1 for ease of comparison, along with % differences from vehicle controls. In both sexes and consistent with previous findings, semaglutide promoted most of the weight loss during the first week, where reductions in body masses were dose-dependent (Figure 1B and 1C, left panels). In females only, 3 nmol/kg semaglutide continued to promote weight loss into the third week, resulting in weight loss that was equivalent between 3 and 30 nmol/kg doses (Figure 1B and 1C, right panels). Altogether, semaglutide promoted 11.3% and 20.2% weight loss in males, and 21.1% and 26.1% weight loss in females (Table 1). On the other hand, food intake did not appear to be robustly different between 3 and 30 nmol/kg doses, except that semaglutide appeared to suppress food intake a few days longer in females compared to males (Figure 1D). In both sexes, food intake returned to vehicle-injected control levels during the second week of semaglutide treatment regardless of the dose.

**Table 1:** Individual muscle and organ tissue weights of male and female aged mice treated with vehicle or semaglutide. Data presented as absolute mass ± SEM and % difference between the groups (n = 6-10/group for male and n = 5-8/group for female mice).

|  | Vehicle |  | 3 nmol |  | 30 nmol |  | Vehicle vs. 3 nmol |  | Vehicle vs. 30 nmol |  |
|  | **Absolute mass ± SEM** |  |  |  |  |  | **% Difference** |  |  |  |
| Sex | Male | Female | Male | Female | Male | Female | Male | Female | Male | Female |
|---|---|---|---|---|---|---|---|---|---|---|
| Body mass (g) | 45.2 ±1.6 | 40.9 ± 1.5 | 40.1 ± 2.2 | 32.3 ± 1.8 | 36 ± 1.8 | 30.2 ± 1.6 | -11.3 | -21.1 | -20.2 | -26.1 |
| Lean mass (g) | 24.9 ± 0.5 | 20 ± 0.4 | 24.3 ± 0.5 | 18.2 ± 0.6 | 23.2 ± 0.6 | 18.3 ± 0.4 | -2.3 | -8.6 | -6.6 | -8.3 |
| Fat mass (g) | 16.7 ± 1.3 | 20.2 ± 1.6 | 16.7 ± 2.0 | 12.0 ± 2.0 | 8.4 ± 1.4 | 8.5 ± 1.7 | -31.8 | -40.6 | -50.1 | -58.0 |
| iWAT (mg) | 1328.3 ± 94 | 1676.7 ± 173.8 | 1072 ± 193.2 | 1226.2 ± 150 | 534.3 ± 96.5 | 516.4 ± 99.2 | -19.3 | -26.9 | -59.8 | -69.2 |
| gWAT (mg) | 2971.6 ± 235 | 2001.1 ± 237 | 2438.4 ± 415 | 1537.9 ± 211.9 | 1769.8 ± 313.2 | 681.9 ± 164.7 | -17.9 | -23.2 | -40.4 | -65.9 |
| TA (mg) | 50 ± 0.9 | 39.8 ± 1.1 | 48 ± 1.7 | 38.4 ± 1.2 | 49.2 ± 1.3 | 38.3 ± 1.7 | -3.9 | -3.5 | -1.4 | -3.7 |
| Gastrocnemius (mg) | 131.4 ± 2.7 | 108.5 ± 2.2 | 126.1 ± 3.5 | 102.5 ± 2.6 | 126.8 ± 4 | 98.3 ± 3.6 | -4.1 | -5.6 | -3.5 | -9.4 |
| Quadriceps (mg) | 194.7 ± 4.2 | 158.3 ± 5.2 | 178.6 ± 8 | 153 ± 5.3 | 183.4 ± 5 | 135.8 ± 6.5 | -8.3 | -3.4 | -5.8 | -14.2 |
| Plantaris (mg) | 17.5 ± 0.4 | 13.5 ± 0.4 | 17.8 ± 0.3 | 13.2 ± 0.3 | 16.9 ± 0.5 | 12.3 ± 0.3 | 1.6 | -2.3 | -3.5 | -8.8 |
| EDL (mg) | 11.8 ± 0.2 | 10.2 ± 0.3 | 11.6 ± 0.4 | 9.7 ± 0.3 | 11.8 ± 0.5 | 9.3 ± 0.3 | -1.5 | -4.8 | 0.5 | -8.7 |
| Soleus (mg) | 9.9 ± 0.4 | 8.0 ± 0.2 | 9.5 ± 0.4 | 7.8 ± 0.5 | 8.9 ± 0.5 | 7.4 ± 0.4 | -3.9 | -3.5 | -10.1 | -8.4 |
| Diaphragm (mg) | 93.2 ± 4 | 83.2 ± 4.4 | 100.2 ± 8.8 | 89.7 ± 7.2 | 88.8 ± 3.1 | 93.0 ± 2.7 | 7.5 | 7.8 | -4.7 | 11.8 |
| Liver (mg) | 2954.9 ± 200.6 | 2119 ± 168.6 | 2184.5 ± 118.2 | 1777.1 ± 137.6 | 1957.5 ± 171.1 | 1702.8 ± 92.1 | -26.07 | -16.13 | -33.76 | -19.64 |
| Spleen (mg) | 92.9 ± 3.2 | 223.6 ± 32.5 | 13.8 ± 29.8 | 211.7 ± 50.8 | 111.2 ± 17.4 | 197.1 ± 35.4 | 40.83 | -5.30 | 19.74 | -11.85 |
| Kidney (mg) | 419.9 ± 12.1 | 365.3 ± 16 | 406.9 ± 19.1 | 371.9 ± 13.6 | 465 ± 31.6 | 430.2 ± 23.9 | -3.10 | 1.81 | 10.73 | 17.76 |
| Heart (mg) | 154.5 ± 4.6 | 134.4 ± 6.6 | 141.1 ± 10.2 | 131.4 ± 8.8 | 142.8 ± 11.7 | 156.5 ± 14.2 | -8.62 | -2.21 | -7.55 | 16.46 |
| Lung (mg) | 189.2 ± 6.4 | 169.5 ± 5.7 | 189 ± 11.0 | 163.8 ± 8.6 | 186.9 ± 7.8 | 190.9 ± 5.7 | 0.17 | -3.34 | -1.20 | 12.64 |

### Aging Attenuates the Loss of Lean Mass Induced by Semaglutide without Attenuating the Loss of Fat Mass

Assessment of body composition by nuclear magnetic resonance (NMR) spectroscopy demonstrated a clear dose-dependent loss of fat mass that were similar in males and females (Figure 2A and 2B, left). This was reflected in similar dose-dependent losses of individual adipose tissue mass in both sexes (Figure 2C and 2D). The loss of fat mass induced by semaglutide was equivalent to those observed in young mice [6, 7, 9, 10].

**Figure 2.**
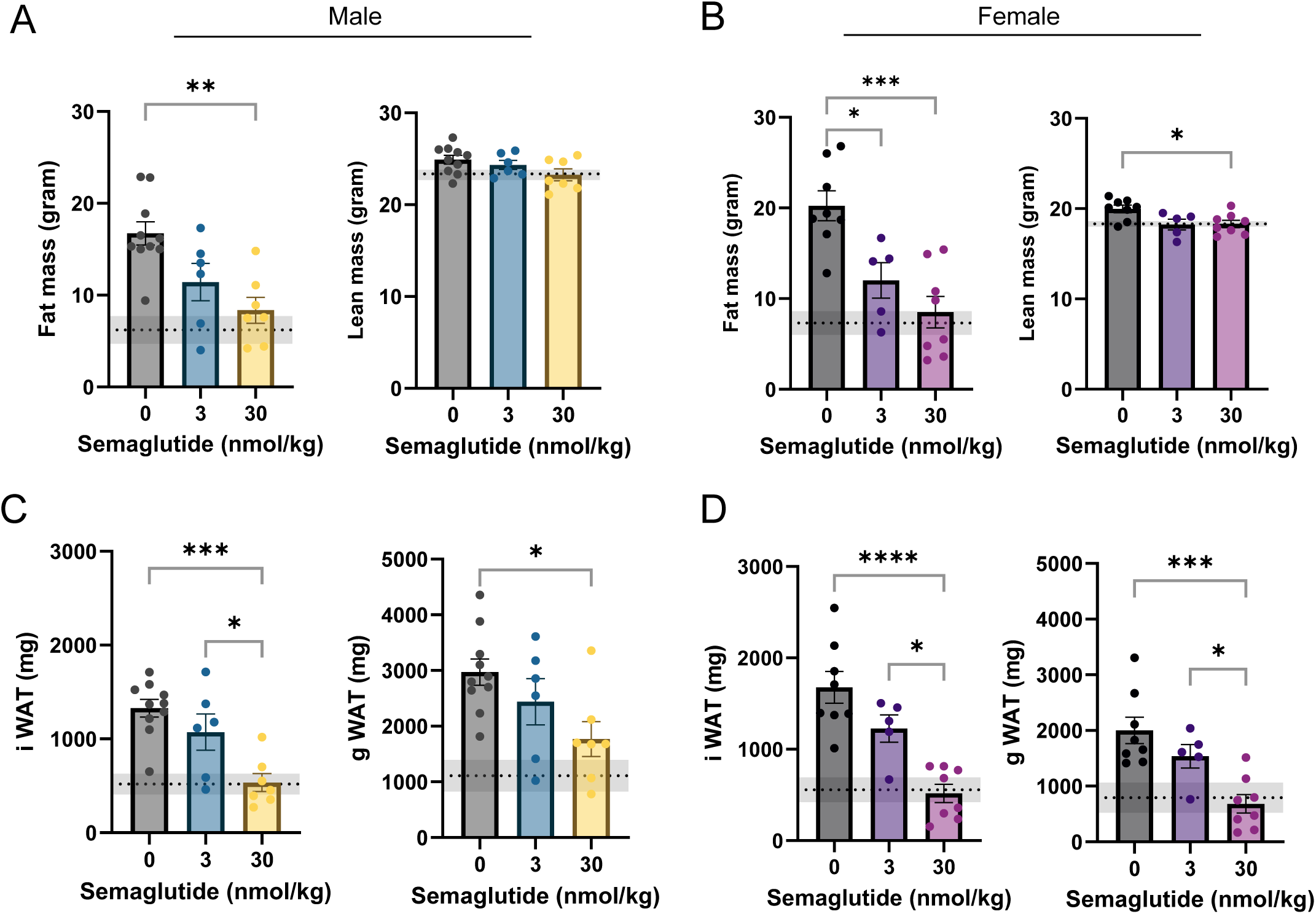
Semaglutide-induced alteration in body composition and fat depots in aged male and female mice. (A) Terminal body composition such as fat mass and lean mass in vehicle and semaglutide treated male mice. (B) Terminal fat and lean mass of vehicle and semaglutide treated female mice. (C) Individual fat depot mass in vehicle or semaglutide treated male mice. (D) Individual fat mass in female mice. All Data is represented as mean ± SEM. * p<0.05, ** p<0.01, *** p<0.001 and **** p<0.0001, significant difference vs. corresponding controls. n = 6-10/group for male and n = 5-8/group for female mice.

In contrast, there appeared to be more nuanced sex differences in the loss of lean mass (Figure 2A and 2B, right), where the effects were dose-dependent in males (2.3% and 6.6%), but not in females (8.6% and 8.3%). Thus, the magnitudes in the loss of lean mass were greater in females compared to males. It is important to note that, regardless of sex, the loss of lean mass induced by semaglutide in aged mice is substantially lower compared to published values for loss of lean mass that occurs in young mice treated with semaglutide [6, 7, 9, 10].

### Aged Female Mice Are More Prone to Semaglutide-induced Loss in Muscle Mass

Central to the premise of the current study, we next examined the effect of semaglutide treatment in skeletal muscle mass. There were striking differences between male and female mice. In males, regardless of doses, semaglutide appeared to have no meaningful effect on five of the limb muscle masses (tibialis anterior, gastrocnemius, quadriceps, plantaris, extensor digitorum longus or EDL) we quantified, with an exception for soleus which had a trend for ∼10% reduction at the higher dose (Figure 3A and Table 1). In contrast, losses of skeletal muscle masses were substantially greater in females particularly with the 30 nmol/kg dose (Figure 3B and Table 1), with masses of the quadriceps 14.2% lower in compared to vehicle control. Indeed, percent losses in five limb muscles were greater than the loss of lean mass (>8.3%) in these mice. This is different from our previous findings in young mice, where loss of individual skeletal muscle masses did not come close to reaching the magnitude of loss in lean mass, as greater proportions of mass loss came from other lean tissues such as liver, spleen, and heart. In aged mice, semaglutide treatment also reduced the liver mass, but did not appear to reduce masses of other central organs (Supplemental Figure 1A and 1B). These findings highlight that, unlike young mice, a greater proportion of loss of lean mass is represented by the loss of skeletal muscle mass in aged mice, particularly in females. The sex-dependent effect of semaglutide treatment in skeletal muscle mass was also reflected in the reduced cross-sectional area in myofibers without robust changes in fiber-type (Figure 3C-3F, Supplemental Figure 2A and 2B).

**Figure 3.**
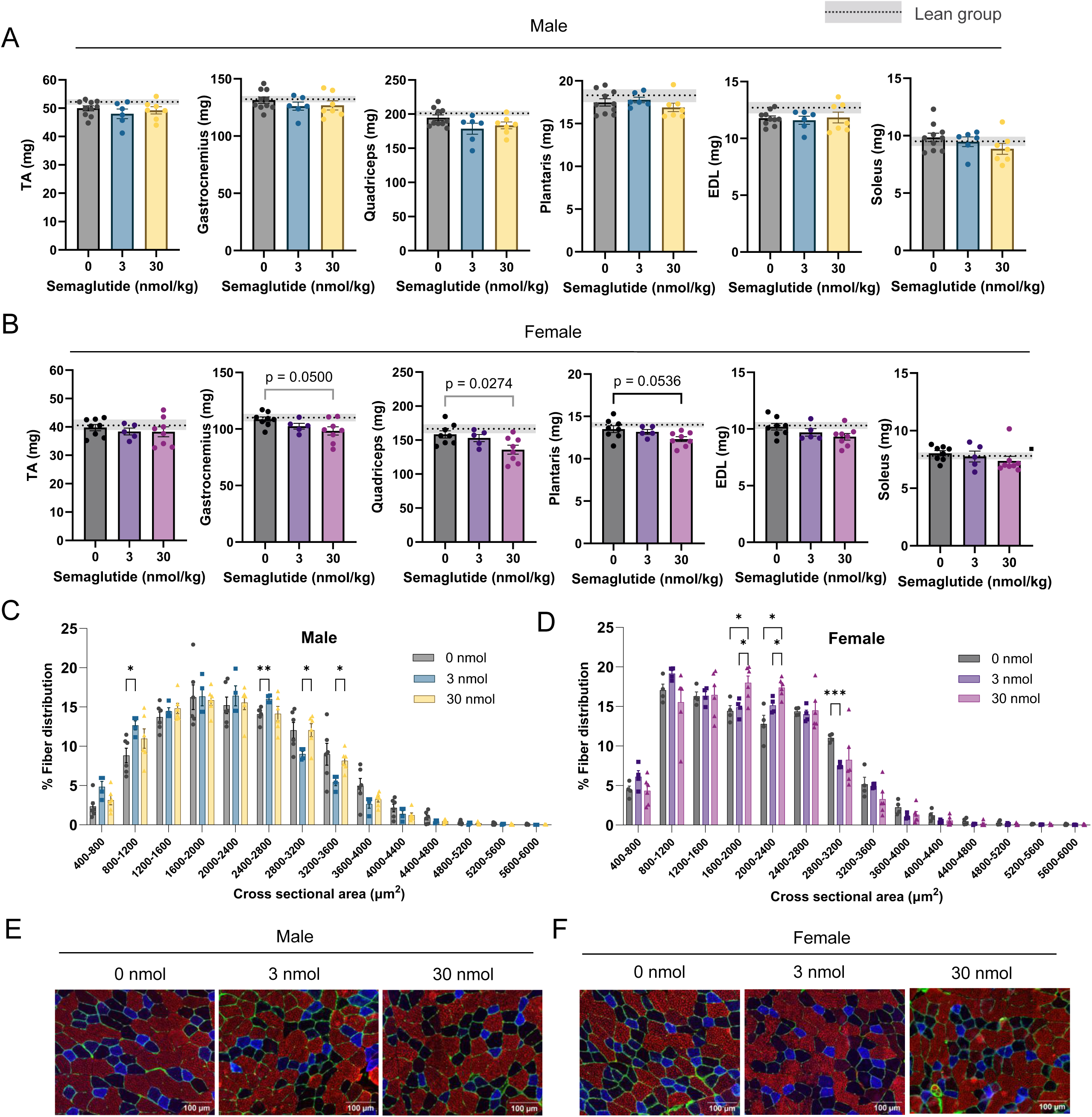
Effects of semaglutide treatment on skeletal muscle mass and cross-sectional area in aged male and female mice. (A) Individual muscle mass of vehicle or 3nmol/kg or 30nmol/kg of semaglutide treatment in male mice. (B) Individual muscle mass of vehicle or semaglutide treated female aged mice. (C and D) Frequency distribution of myofiber cross sectional area in male and female mice respectively. (D and E) Immunohistochemical staining showing different muscle fibers (red-Type IIb, blue-Type IIa and black-Type IIx) in both male and female mice respectively. All Data is represented as mean ± SEM. * p<0.05 significant difference vs. corresponding controls. n = 6-10/group for male and n = 5-8/group for female mice and n = 4-6/group/sex for immunohistochemical staining.

Another important difference in observations between young and aged mice emerge when comparing muscle masses to age-matched lean controls (Figure 3A and 3B, dotted lines with grey zones). In young mice, HFD feeding increased masses of all limb skeletal muscles [6]. Greater weight loading and anabolism due to obesity and/or high caloric intake likely contributed to these increases in skeletal muscle mass. Thus, in young mice, a concern for the small losses in muscle masses was attenuated by the observations that muscle masses after semaglutide treatment remained essentially equivalent to those in lean mice. In contrast, HFD feeding very clearly did not increase masses of limb skeletal muscle in aged male or female mice (Figure 3A and 3B). Thus, muscle masses in semaglutide-treated mice were not only lower compared to vehicle control, but also from age-matched lean control.

### Aged Male Mice Are More Prone to Semaglutide-induced Loss in Muscle Strength

In young mice reported in our previous article, one of the main findings was that semaglutide-treatment induced a dose-dependent decrease in skeletal muscle force-generating capacity that was disproportionately more compared to the small loss in skeletal muscle mass [6]. Nevertheless, HFD feeding also induced increases in force-generating capacity, such that semaglutide treatment only brought them down to the level of the age-matched lean mice. In contrast, in aged mice in the current study, HFD feeding only had a trend for a small increase in force-generating capacity in soleus muscle, and no trend in muscle EDL (Figure 4A and 4B, compared to dotted lean control). Thus, similar to our findings in muscle mass (Figure 3), any loss in force induced by semaglutide would also mean loss of strength below age-matched lean mice.

**Figure 4:**
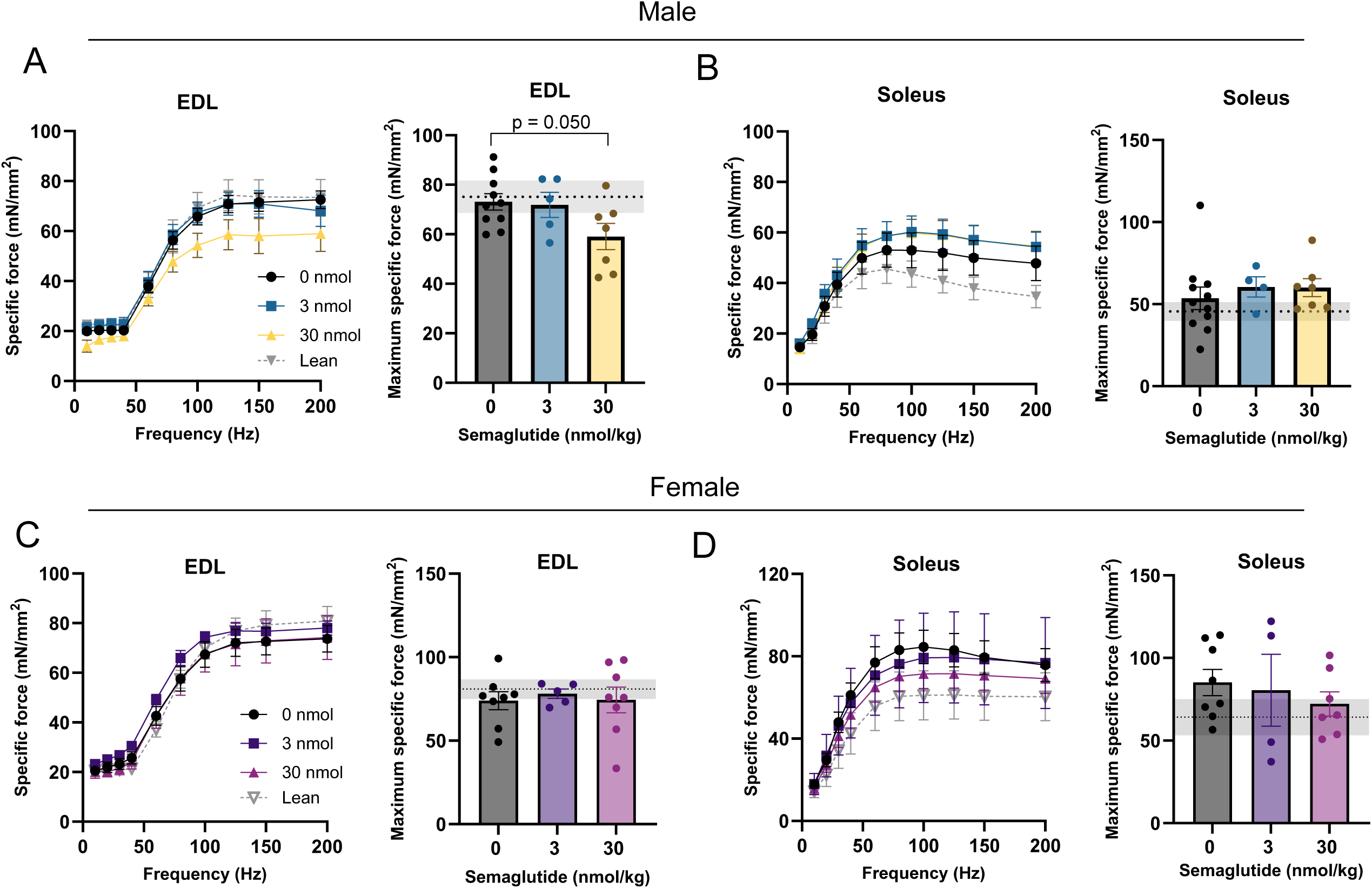
Effects of semaglutide on skeletal muscle force-generating capacity in aged male and female mice. (A) Specific force production and maximal force-generating capacity of EDL muscle with vehicle or semaglutide treatment in aged male mice. (B) Specific force production and maximal force-generating capacity of soleus muscle in aged male mice. (C) Specific muscle force production and maximal force-generating capacity of EDL muscle in aged female mice. (D) Specific muscle force production and maximal force-generating capacity of soleus muscle in aged female mice. All Data is represented as mean ± SEM. * p<0.05 significant difference vs. corresponding controls. n = 6-10/group for male and n = 5-8/group for female mice.

In aged males, only with the 30 nmol/kg dose, there was a substantial decrease (19.2%) in maximal force produced by EDL muscle (Figure 4A and Supplemental Figure 3A). It is interesting to note that EDL muscle did not undergo atrophy under the same condition in aged male mice (Figure 3A). Semaglutide did not influence force-generating capacity in soleus muscles in aged male mice (Figure 4B and Supplemental Figure 3B). In aged females, semaglutide treatment had no effect on skeletal muscle force generating capacity in soleus or EDL muscle (Figure 4C and 4D and Supplemental Figure 3C and 3D).

## Conclusion

In summary, our findings suggest that aging may augment concerns associated with semaglutide-induced weight loss on skeletal muscle. In aged female mice, 30 nmol/kg of semaglutide induced reductions in the masses of hindlimb muscles (14.2% in quadriceps and 9.4% in gastrocnemius, the largest two muscles we examined) that exceeded the percent reduction in the whole-body muscle mass (8.3%). In aged male mice, the high dose semaglutide reduced the maximal force production of EDL muscle by 19.2%. In both cases, the concern is amplified by our observations that, unlike in our studies in young mice that used identical study design [6], HFD feeding did not increase muscle mass or force in aged mice. These observations raise questions regarding the potential consequences of pharmacologically induced weight loss for skeletal muscle mass and strength in older populations, men or women.

Given the limitations of translating findings from mice to humans, clinical studies are essential to determine whether similar age- and sex-specific responses occur in individuals receiving incretin-based therapies. It is noteworthy that FDA has now approved a higher dose of semaglutide treatment (7.2 mg/week) [16], which is thought to be approximately equivalent to the 30 nmol/kg/day dose of semaglutide treatment in mice for producing similar magnitude of weight loss. Ongoing clinical trials evaluating skeletal muscle mass and function using validated measurement techniques should prioritize the inclusion of older adults, particularly those at increased risk of sarcopenia, to better characterize the long-term implications of GLP-1RA-induced weight loss for muscle health.

## Acknowledgment

This research is supported by NIH grants (DK107397, DK127979, and AG074535 to K.F), and the Grant-in-aid for Japan, Society for Promotion of Science (JSPS) Fellows (24KJ2039 to T.K.).

## Ethics statement

All animal experiments were performed with the approval of the Institutional Animal Care and Use Committees at the University of Utah.

## Supplemental Figure Legends

**Supplemental Figure 1:**
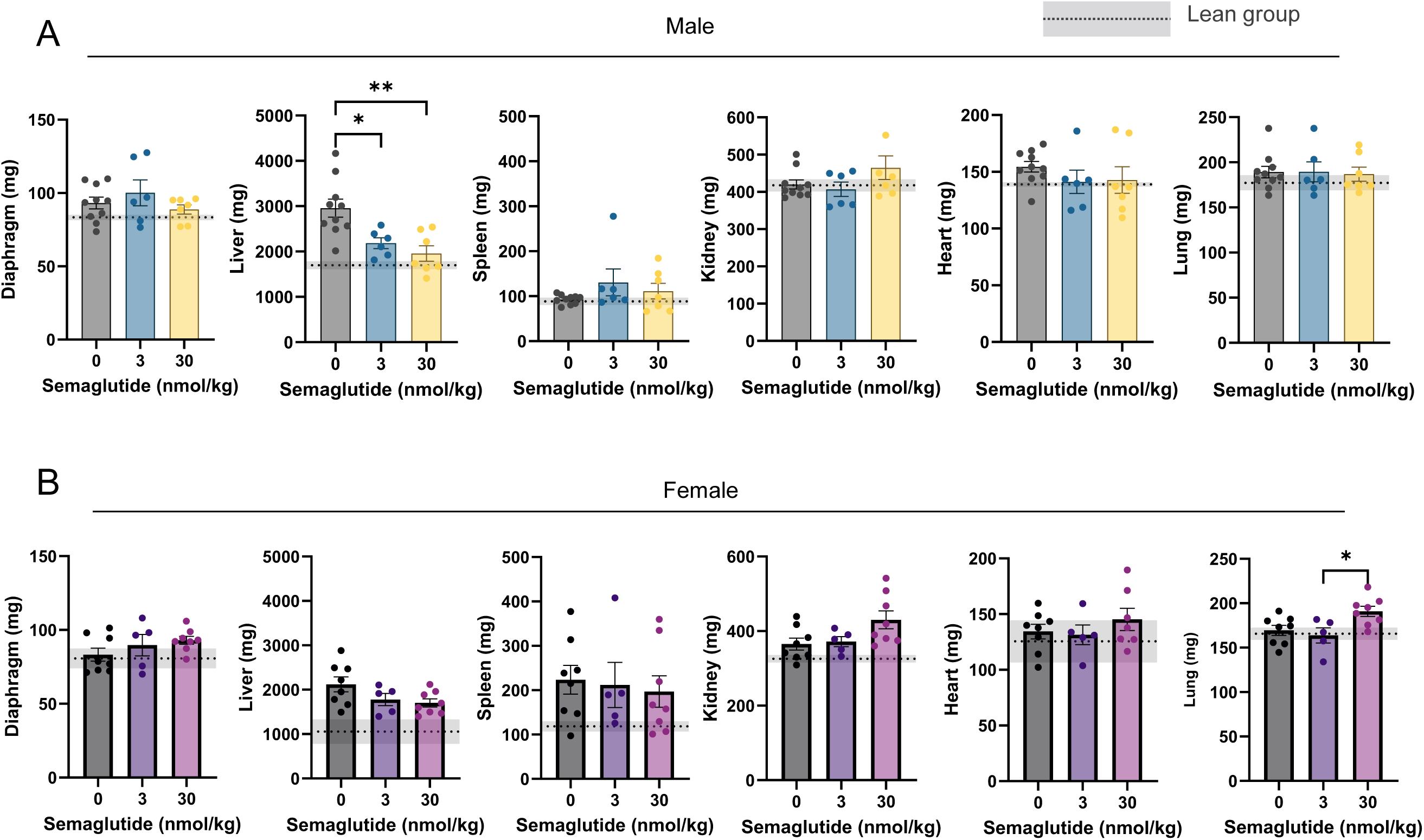
Effects of semaglutide treatment on individual organ mass and in aged male and female mice. (A) Individual organ mass in vehicle or 3nmol/kg or 30nmol/kg of semaglutide treatment in male mice. (B) Individual organ mass in vehicle or 3nmol/kg or 30nmol/kg of semaglutide treatment in female mice. All Data is represented as mean ± SEM. * p<0.05, and ** p<0.01 significant difference vs. corresponding controls. n = 6-10/group for male and n = 5-8/group for female mice.

**Supplemental Figure 2:**
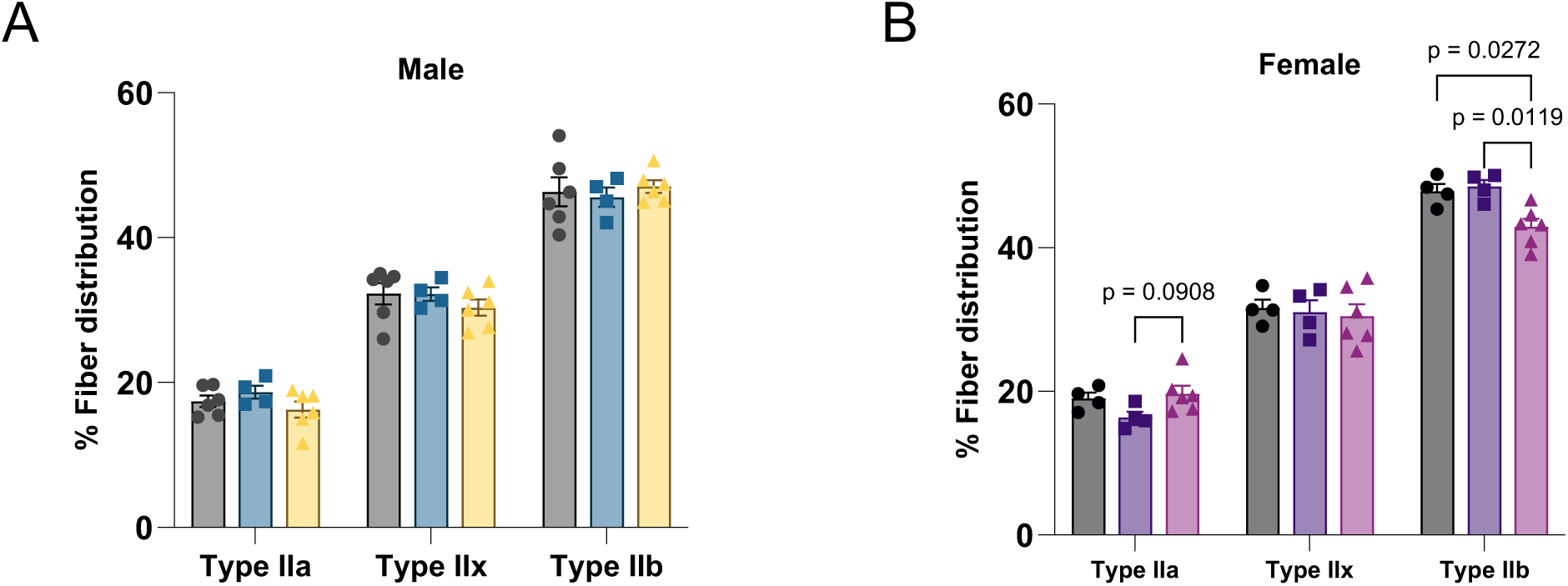
Effects of semaglutide on skeletal muscle fiber types in aged male and female mice. (A) Percentage of muscle fiber type distribution analysis from immunohistochemical staining in vehicle or semaglutide treated both male mice. (A) Percentage of muscle fiber type distribution analysis from immunohistochemical staining in vehicle or semaglutide treated both female mice. All Data is represented as mean ± SEM. * p<0.05, and ** p<0.01 significant difference vs. corresponding controls. n = 4-6/group/sex for immunohistochemical staining.

**Supplemental Figure 3:**
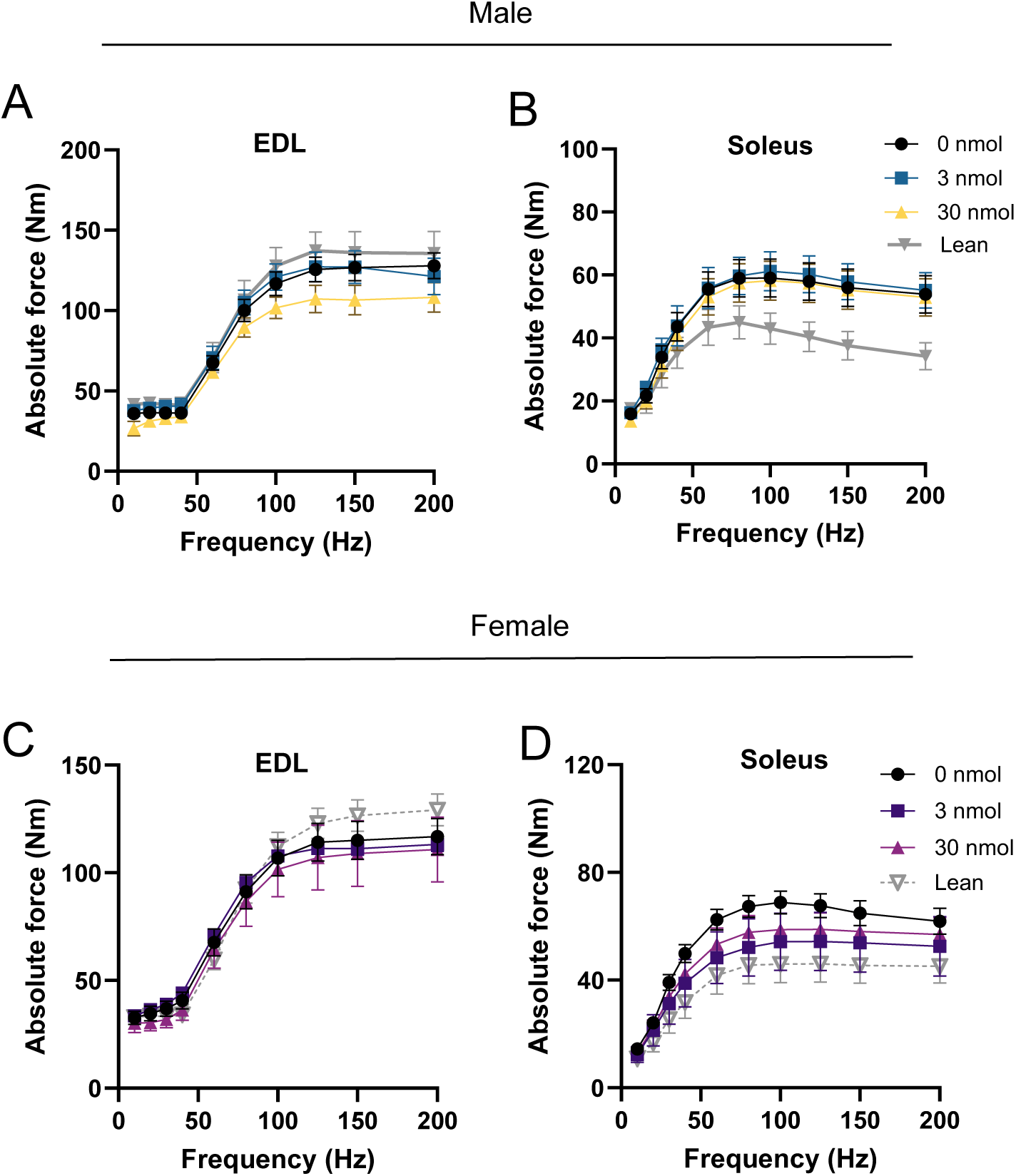
Effects of semaglutide on skeletal muscle force-generating capacity in aged male and female mice. (A) Absolute force production of EDL muscle with vehicle or semaglutide treatment in aged male mice. (B) Absolute force production of soleus muscle in aged male mice. (C) Absolute force production of EDL muscle in aged female mice. (D) Absolute force production of soleus muscle in aged female mice. All Data is represented as mean ± SEM. * p<0.05 significant difference vs. corresponding controls. n = 6-10/group for male and n = 5-8/group for female mice.

## References

1. Wilding JPH, Batterham RL, Calanna S, Davies M, Gaal LFV, Lingvay I, et al. Once-Weekly Semaglutide in Adults with Overweight or Obesity. New England Journal of Medicine. 2021;384:989–1002. doi:doi:10.1056/NEJMoa2032183

2. Bergmann NC, Davies MJ, Lingvay I, Knop FK. Semaglutide for the treatment of overweight and obesity: A review. Diabetes Obes Metab. 2023;25:18–35. doi:10.1111/dom.14863

3. Wilding John PH, Batterham Rachel L, Calanna S, Davies M, Van Gaal Luc F, Lingvay I, et al. Once-Weekly Semaglutide in Adults with Overweight or Obesity. New England Journal of Medicine. 2021;384:989–1002. doi:10.1056/NEJMoa2032183

4. Neeland IJ, Linge J, Birkenfeld AL. Changes in lean body mass with glucagon-like peptide-1-based therapies and mitigation strategies. Diabetes Obes Metab. 2024;26 Suppl 4:16–27. doi:10.1111/dom.15728

5. Tinsley GM, Heymsfield SB. Fundamental Body Composition Principles Provide Context for Fat-Free and Skeletal Muscle Loss With GLP-1 RA Treatments. J Endocr Soc. 2024;8:bvae164. doi:10.1210/jendso/bvae164

6. Karasawa T, Choi RH, Meza CA, Rout S, Drummond MJ, Chaix A, et al. Unexpected effects of semaglutide on skeletal muscle mass and force-generating capacity in mice. Cell Metab. 2025;37:1619–20. doi:10.1016/j.cmet.2025.07.004

7. Langer HT, Gilmore NK, Hayden CMT, Roux J, Bariohay B, Rouquet T, et al. Weight loss with GLP-1 medicines does not result in a disproportionate loss of muscle mass or function in obese mice and humans. Cell Reports Medicine. 2026;7:doi:10.1016/j.xcrm.2026.102665

8. Rout S, Karasawa T, Watanabe S, Chaix A, Drummond MJ, Funai K, et al. Females Are Completely Resistant to Semaglutide-Induced Muscle Loss in ob/ob Mice. Diabetes. 2026;75:1584–95. doi:10.2337/db26-0229

9. Nunn E, Jaiswal N, Gavin M, Uehara K, Stefkovich M, Drareni K, et al. Antibody blockade of activin type II receptors preserves skeletal muscle mass and enhances fat loss during GLP-1 receptor agonism. Mol Metab. 2024;80:101880. doi:10.1016/j.molmet.2024.101880

10. Jeromson S, Baranowski B, Akcan M, Waters BD, Eisner K, Bellucci A, et al. Semaglutide impacts skeletal muscle to a similar extent as caloric restriction in mice with diet-induced obesity. J Physiol. 2025;n/a:doi:10.1113/JP289449

11. Abuetabh Y, Schmidt MA, Naganuma M, Vaka R, El-Ghiaty MA, Braun S, et al. Semaglutide-induced loss of skeletal muscle mass is blunted by co-administration of ketone esters. JCI Insight. 2026;11:doi:10.1172/jci.insight.201810

12. Fryar CD, Afful J, Saif NT. Prevalence of Overweight, Obesity, and Severe Obesity Among Adults Age 20 and Older: United States, 1960-1962 Through August 2021-August 2023. In: Hyattsville (MD): 2024.

13. Reidy PT, McKenzie AI, Mahmassani ZS, Petrocelli JJ, Nelson DB, Lindsay CC, et al. Aging impairs mouse skeletal muscle macrophage polarization and muscle-specific abundance during recovery from disuse. American Journal of Physiology-Endocrinology and Metabolism. 2019;317:E85–E98. doi:10.1152/ajpendo.00422.2018

14. Vahratian A, Warren A. GLP-1 Injectable Use Among Adults With Diagnosed Diabetes: United States, 2024. NCHS Data Brief. 2025;doi:10.15620/cdc/174616

15. Yanai S, Endo S. Functional Aging in Male C57BL/6J Mice Across the Life-Span: A Systematic Behavioral Analysis of Motor, Emotional, and Memory Function to Define an Aging Phenotype. Front Aging Neurosci. 2021;13:697621. doi:10.3389/fnagi.2021.697621

16. Wharton S, Freitas P, Hjelmesaeth J, Kabisch M, Kandler K, Lingvay I, et al. Once-weekly semaglutide 7.2 mg in adults with obesity (STEP UP): a randomised, controlled, phase 3b trial. Lancet Diabetes Endocrinol. 2025;13:949–63. doi:10.1016/S2213-8587(25)00226-8

